# Neretva: Neural Variational Inference for Allele-level Genotyping of Highly Polymorphic Genes

**DOI:** 10.64898/2026.02.03.703582

**Authors:** Qinghui Zhou, Seyed Pouria Ahmadi, Ibrahim Numanagić

## Abstract

Accurate genotyping and phasing of highly polymorphic gene families are essential for precision medicine. Yet, the genotyping problem remains computationally challenging due to extreme sequence similarity between related genes, copy number variation, and structural complexity. Current methods typically rely on integer linear programming or maximum likelihood-based approaches that often suffer from scalability and flexibility issues. Here, we present Neretva, a new framework that models the genotyping problem as a probabilistic latent variable model and employs auto-encoding variational Bayes (AEVB) for inference. Leveraging neural inference networks, our approach scales efficiently to complex gene families such as CYP (cytochrome P450) and KIR (killer-cell immunoglobulin-like receptors), and achieves competitive or improved accuracy over the current state of the art on both families. Neretva is open source and is freely available at https://github.com/0xTCG/neretva.

## Introduction

High-throughput sequencing (HTS) technologies are a promising solution for enabling precision medicine through comprehensive and rapid analysis of genotypes of clinically important gene families [48]. Key families include the cytochrome P450 (CYP) pharmacogenes and killer-cell immunoglobulin-like receptor (KIR) genes. The CYP family encodes enzymes responsible for the metabolism of more than 70% of clinically prescribed drugs [49], while KIR genes regulate natural killer cell and T-cell responses, and are linked, among other things, to HIV progression, autoimmune disorders, and organ transplant success rates [30, 1, 32].

Despite its importance, accurate genotyping of these genes in HTS data remains challenging due to several factors. The first challenge stems from the fact that their alleles are defined as haplotypes, which requires accurate phasing to establish a clinically actionable phenotype. Phasing is often hindered by short read length, as well as extreme inter-gene sequence similarity between pharmacogenes and their paralogs (often present within pharmacogene clusters [36]) and the intragene allelic identity (often exceeding *>*98%), which introduce significant ambiguity during assigning individual HTS reads to their true origin [4, 35]. Many pharmacogene families also exhibit substantial population-specific variation [47], thus rendering the strict reliance on the previously characterized alleles insufficient. Furthermore, some pharmacogenes exhibit extensive copy number variation and are subject to complex structural variations such as gene deletions, duplications, and paralog-related gene fusions [12]. Clusters such as KIR exhibit even higher levels of complexity; around half of the 17 KIR genes exhibit extensive diversity and are subject to copy number variation and cross-gene structural rearrangements [17, 18]. Finally, we note that other clinically important gene families not covered here present similar challenges; examples include *HLA* genes, immunoglobulin loci, T-cell receptor genes, and other immune receptor families such as *FCGR* and *LILR* [10, 2, 28, 43].

Many specialized frameworks were developed to accurately infer such genotypes from HTS data. Tools such as Stargazer [25], Cyrius [7], StellarPGx [45], PyPGx [26], Aldy [35, 15] and Astrolabe [46] combine the existing allele databases such as PharmVar [13], as well rule-based strategies and various statistical and combinatorial techniques, to accurately infer genotypes for wide variety of pharmacogenes. For genotyping KIR genes, such as T1K [41], Locityper [38], Geny [50], kir-mapper [5] and PING [31] often rely on combinatorial and likelihood-based assignment strategies to maximize the likelihood of observed read data and estimate candidate allele abundances, together with various pre-processing schemes such as *k*-mer-based fingerprinting. Other gene clusters are also handled by specialized tools; some examples include OptiType [43] and HLAreporter [16] (for HLA), Immunotyper-SR (for IGH) [10, 11], and the recent pangenome-based approaches such as PGR-TK [8].

As demonstrated by the performance of the tools such as Aldy (pharmacogenes)^1^, Geny (KIR), ImmunoTyper-SR (IGH), OptiType (HLA), as well as the recent pangenome-based PHI [6], one of the most effective ways to genotype complex genes is through combinatorial optimization [34, 44]. In this approach, the candidate alleles are first filtered by eliminating alleles with low abundance (through a rule or statistical-based approach), and the optimal solution is then found by solving one or more constrained integer linear programming (ILP) models, often composed into a multi-stage process where each stage handles a different aspect of the genotyping problem (e.g., Aldy employs separate ILPs for structural variant detection and for allele phasing). Although such approaches guarantee a theoretical optimum and constraint enforcement, and avoid converging to an incorrect local optimum (unlike with likelihood-based approaches), they are often NP-hard [35] and scale poorly to new data types and complex gene families. While such models are feasible and practical for pharmacogenes (where the ILP instances contain a manageable number of variables), handling of complex gene families such as KIR leads to a combinatorial explosion that results in either impractical runtimes or in sacrificing optimality guarantees. ILP-based approaches also rely on black-box (and often proprietary) solvers such as Gurobi or Google OR-Tools, which make their use impractical in specialized clinical applications that require use of privacy-preserving technologies (e.g., homomorphic encryption) for handling patient data. Lastly, ILP solutions are rigid, and it is not easy to establish the confidence of an ILP solution in a noisy dataset.

These limitations motivate a probabilistic ILP relaxation that can enable scalable gradient-based optimization while naturally accommodating uncertainty. Recent advances in variational inference, such as *auto-encoding variational Bayes* (AEVB) have shown remarkable success in high-dimensional problems with a similar latent structure to the genotyping problem. In natural language processing, one such example is NMF-fashioned LDA [24, 3], while neural variational models such as ProdLDA [42], NVDM [33] and ETM [9] have scaled topic modeling to millions of documents and thousands of topics. In genomics, deep learning approaches such as CAECseq [20] (which uses convolutional autoencoders for haplotype assembly and viral quasispecies reconstruction) have demonstrated that neural networks can effectively cluster sequencing reads originating from highly similar genomic regions. All these methods leverage GPU-accelerated gradient optimization and inference networks to achieve orders-of-magnitude speedup over traditional approaches without impacting the model accuracy. Beyond scalability, AEVB models also offer flexibility for incorporating domain constraints as differentiable regularizers, and avoid the need to re-derive update equations (as in likelihood-based strategies) or redesign samplers (as in MCMC), which makes them desirable for large-scale inference problems like KIR genotyping.

In this paper, we present a new unified AEVB-based framework, called Neretva, for genotyping both pharmacogene (CYP) and KIR gene families. Our approach combines the variant-centric modelling philosophy of Aldy with the scalability of modern variational inference. We demonstrate that Neretva achieves high accuracy, precision and recall across both gene families, with *F*_1_ scores reaching up to 1.0 in CYP and 0.912 in KIR experiments, thus matching or surpassing the current state of the art while maintaining the interpretability of the results essential for precision medicine applications. Thus, we hope that Neretva will serve as a sound and practical solution for employing HTS data to genotype complex pharmacogenes in both research and clinical settings.

## Methods

Given a complex gene family and a collection of sequencing reads in BAM/CRAM/FASTQ, Neretva infers the overall genotype of the given family by (1) determining the number of copies of each gene within that family, and by (2) constructing the exact sequence content (haplotype) of each such copy (in other words, assign an allele designation to it).

### Database Preparation

Consider a multi-gene model where 𝒢 = {*G*_1_, …, *G*_*L*_} represents a family of *L* ≥ 1 genes (e.g., KIR family contains *L* = 17 genes). Let us define a set ℳ_*G*_ of all possible variants occurring within *G* as 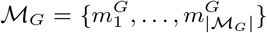, and partition it into 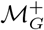 consisting of *core variants* (that impact the phenotype or a protein function of *G*), and 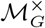 that contain *minor variants* (silent variants without known functionality).

Each gene *G* ∈ 𝒢 defines a *reference allele* with no variants (often referred to as *1 allele). Each other allele 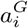 of that gene is defined by a set of variants (denoted as 𝒜^*G*^), itself split into a set of core 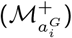 and minor 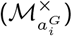 variants, that differentiates such allele from the reference allele. We will later refer to 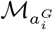 as allele 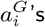 *original variant set*, and define ℳ = ⋃_*G*∈𝒢_ ℳ_*G*_ as a set of all variants being considered.

For successful genotyping, it is essential to construct a database of previously characterized alleles that includes the known alleles for each gene of interest, along with the variant and structural variation information. For KIR genes, we rely on IPD-KIR database v2.12.0 [40], while for other genes, we utilize the PharmVar database 6.2 [13]. In both databases, allele names roughly follow the star-allele nomenclature [39], where the major allele designation (typically represented as star-number; e.g., *CYP2D6* 2) describes the allele’s functionality (and is defined by its core variants), while the minor allele designation (e.g., *2A) distinguishes alleles with differing minor variations.

### Alignment and Filtering

Consider an HTS sample with reads ℛ = {*r*_1_, …, *r*_|ℛ|_}. For pharmacogenes, we assume each read alignment to be correct and use it as-is. As for KIR, where many reads ambiguously map to different genes, we align reads against all known allele sequences with minimap2 [27] in all-to-all mode^2^. Then, for each *r* ∈ ℛ, we gather a set of alleles 𝒜_*r*_ to which *r* maps, and a set of variants ℳ_*r*_ ⊆ ℳ that *r* supports. To reduce the search space, we filter down the allele set 𝒜 into 𝒜^′^ so that each *A* ∈ 𝒜^′^ has each 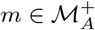 covered by at least 3 reads.

In clusters where incorrect cross-gene alignments are common (e.g., KIR), it is not clear if an observed variant 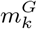 is indeed a true variant in *G* or a *shadow variant* caused by a mapping artifact. An incorrect variant assignment can result in an incorrect allele call, which, in turn, can lead to improper medical treatment. Hence, for each read *r* that induces a variant 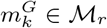, we check whether there is an allele 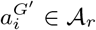 such that *G*^′ ≠^ *G*. If so, we extend 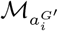 to include 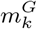 as well. This allows us to construct an *extended variant set* 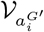 for each allele 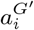, which contains both true *G*^′^ variants and the shadow variants that stem from incorrect alignments. This “extension” has no biological implications; it is only used to guide subsequent optimization steps towards the correct solution.

### Allele Identification Problem

Let us define, for each gene *G*, a list of positions 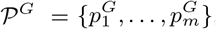 that describe genomic coordinates which can harbor a mutation 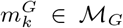 and the set of all positions as 𝒫 = ⋃_*G*∈𝒢_ 𝒫 ^*G*^. We also define a set of *bases* ℬ = {*A, C, T, G, N, P* }, where *A, C, T, G* stand for the four possible nucleotides, while *N* and *P* denote deletions and insertions, respectively. (Note that base is not the same as variant; the reference allele still contains bases despite having no variants.)

After the alignment, we observe a base count matrix **Y** ∈ ℕ^|𝒫|×|ℬ|^ collected from the read set ℛ, where each entry **Y**_*p,b*_ denotes the observed count for base *b* ∈ ℬ at position *p* ∈ 𝒫:

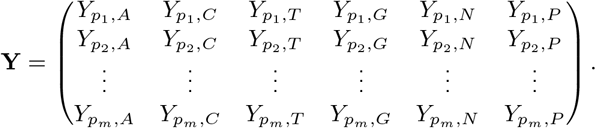

Note that we cannot assume that the allele database can solely explain **Y** due to the possibility of novel (i.e., uncataloged) alleles in a given sample [51]. Thus, to identify the set of alleles that best describes **Y**, we must determine the respective base for each 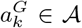 at each *p* ∈ 𝒫 (if *p* describes any variant in 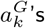 *extended variant set*). Both the allele proportions and the base emissions are modelled as latent variables; more precisely, ***θ*** ∈ Δ^*K*^ represents the relative proportion of each of *K* = |𝒜^′^| candidate alleles, where Δ^*K*^ denotes a *K* − 1 simplex. **Ψ** = {*ψ*_*a,p*_ | *a* ∈ 𝒜^′^, *p* ∈ 𝒫} represents the base emission probabilities, where each *ψ*_*a,p*_ ∈ Δ^|ℬ|^ is a distribution over bases ℬ at position *p* for an allele *a*.

### Copy Number Estimation

The allele identification determines relative allele proportions ***θ*** on the simplex, thus revealing which alleles are present, as well as their relative abundances. However, before determining an exact count of each allele of interest, we first need to estimate the underlying gene copy numbers, as alleles are constrained by the copy number of their parent genes.

Let us formulate the gene copy number estimation as a coverage reconstruction problem. We observe aggregated coverage values across *C* genomic regions (typically representing introns and exons), where each region’s coverage *c*_*r*_ represents the mean read depth normalized by the expected coverage depth. The observed coverage is modelled as a linear combination of unknown gene copy numbers. Let **c** ∈ ℝ^*C*^ denote the observed region coverage vector, **S** ∈ ℝ^*L*×*C*^ denote the gene-to-region contribution matrix where element *S*_*g,r*_ represents the expected coverage contribution rate from gene *G*_*g*_ to region *r*. Let **n** ∈ ℝ^*L*^ denote the unknown copy numbers for *L* genes such that 0 ≤ *n*_*g*_ ≤ *n*_max_. For KIR genes, we set *n*_max_ = 5, while for pharmacogenes, we use *n*_max_ = 4. Then, the expected coverage for each region is given by:

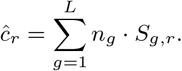

We estimate copy numbers via linear regression by minimizing the following error:

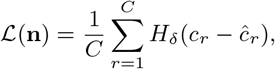

where *H*_*δ*_ is the Huber loss with threshold *δ* that provides robustness to outliers from coverage anomalies. The final copy number is obtained by minimizing ℒ(**n**) via Adam optimizer [21].

### Bias Factor Matrix

While many HTS experiments in theory promise uniform coverage, coverage of some bases might not match the expected coverage due to sequencing biases or mapping artifacts, especially in complex families such as KIR. To model such biases, we define a *bias matrix* **H** ∈ ℝ^*K*×*M*^ for each *a* ∈ 𝒜^′^ and *v* ∈ 𝒱_*a*_ (where *M* = |𝒫| × |ℬ|, *K* = |𝒜^′^|). Its each element *h*_*a,m*_ represents the expected coverage for a variant *m* ∈ ℳ_*a*_.

For variants that are part of an allele’s original variant set, the bias factor is initialized to single copy coverage, or, when available, derived from a predetermined gene profile that accounts for known coverage biases. For shadow variants, we assign bias factors based on empirical misalignment rates obtained from the simulated alignment data. Such matrix is sparse because it has no entries for positions where no variant (true or shadow) is observed. This bias adjustment allows the model to appropriately amplify the signal from the true variants.

### Allele Identification Model

Alleles are inferred by a generative process for the count matrix **Y** collected from a data sample. Let ℒ𝒩(*µ*, Σ) be a logistic normal distribution. For clarity, we define 𝒮 as the set of all allele-position pairs such that 𝒮 = {(*a, p*) | *a* ∈ 𝒜′, *p* ∈ 𝒫}, where each element *s* ∈ 𝒮 represents a specific allele-position pair. The generation process is as follows:

1. draw allele proportions ***θ*** ∼ ℒ𝒩(**0, I**);
2. for each allele position *s* ∈ 𝒮, draw base emissions ***ψ***_*s*_ ∼ ℒ𝒩(***µ***_*s*_, **Σ**_*s*_);
3. for the observed mutation counts matrix, (a) compute mutation probabilities ***π*** ∝ ***θ***^*T*^ ***B***_*η*_, and (b) draw counts **Y** ∼ Multinomial(*N*, ***π***).

Here, **Σ**_*s*_ is assumed to be a diagonal covariance matrix, and **B** ∈ ℝ^*K*×*M*^ denotes the matrix of all the allele base emission probability vectors, **H** ∈ ℝ^*K*×*M*^ is the bias matrix containing the bias factors, and **B**_*η*_ = **B** ⊙ **H** is the strength-adjusted emission matrix. *N* denotes the total observed variant counts across all positions 𝒫 and *N* = Σ_*p,b*∈**Y**_ *Y*_*p,b*_. The database-informed priors ***µ***_*s*_ encode confidence in known variants from the reference database. Under this generative procedure, the marginal likelihood of the observed data is:

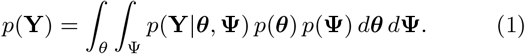

### Variational Inference

The marginal likelihood in Equation (1) is intractable due to the high-dimensional latent space. Thus, we employ variational inference with the SGVB estimator [22] to approximate the posterior *p*(***θ*, Ψ**|**Y**) for each sample independently. For computational efficiency, we adopt mean-field variational approximation [19]:

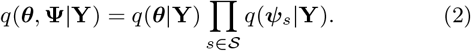

Under this factorization, the evidence lower bound (ELBO) becomes (see Appendix for the complete derivation):

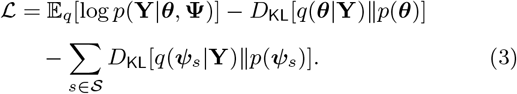

However, while maximizing (3) produces a valid data fit across all variants, it does not prioritize core variants that capture the true biological phenotype of each allele. To address this, consider a set of all core variants 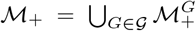 over all genes *G* ∈ 𝒢. Now, consider two probability vectors over ℳ_+_: (1) the empirical functional variant count distribution 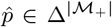 derived from **Y**, and (2) the reconstructed functional variant count distribution 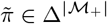 derived from our inference model. We can measure the discrepancy between 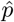 and 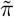 via the Jensen–Shannon divergence [29], whose symmetry allows us to penalize both the lack and the surplus of core variants.

Denote 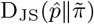 as the Jensen–Shannon divergence between empirical 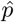 and the estimated core variant count distribution 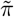. We also regularize the base emission distributions **Ψ** with an entropy penalty. Since each allele possesses a single true base at each genomic position and sequencing error rates are low (typically less than 1% for Illumina platforms), we expect **Ψ** to be sharply peaked rather than diffused. We use an entropy penalty *H*(**Ψ**) = 𝔼_*s*∈𝒮_[*H*(***ψ***_***s***_)] to encourage the model to commit to a dominant base at each position, and acquire the augmented loss term as:

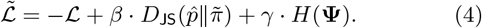

The hyperparameters *β, γ* are balancing terms for the final loss term. We minimize this loss with Adam optimizer [21], where gradients are computed via the reparameterization trick [22] (see Appendix for details).

## Results

We chose three pharmacogene families in the increasing order of complexity to comprehensively benchmark Neretva ^3^. The first family, CYP2C, consists of three genes (*CYP2C8, CYP2C9*, and *CYP2C19*) whose alleles are defined solely in terms of variants, and which do not exhibit any copy number changes, structural variants or mapping ambiguities. The other family, CYP2D, consists of *CYP2D6* and its paralog *CYP2D7*, and exhibits both copy number changes and structural variants. Finally, the most complex KIR family contains 17 genes and, in addition to copy number and structural variants, exhibits high levels of ambiguous mappings.

The pharmacogene (CYP) families are evaluated on 70 GeT-RM Illumina WGS samples with an average sequencing depth of 30× [15]^4^. All ground truths were previously validated by the GeT-RM consortium and augmented by the recently published updates [14, 37, 15]. This dataset contains, in addition to diploid genotypes, a number of *CYP2D6* copy number variants (both deletions and duplications) and paralog-related fusions. Besides Neretva, we also benchmarked the current state-of-the-art pharmacogene genotypers: StellarPGX, Aldy, Cyrius, Stargazer, Astrolabe and PyPGx. Note that, despite the fact that CYP pharmacogene genotyping is a well-studied problem (most tools routinely obtain a near-perfect *F*_1_ score), we still benchmark it to show that our model meets the expected baseline performance.

**Figure 1.**
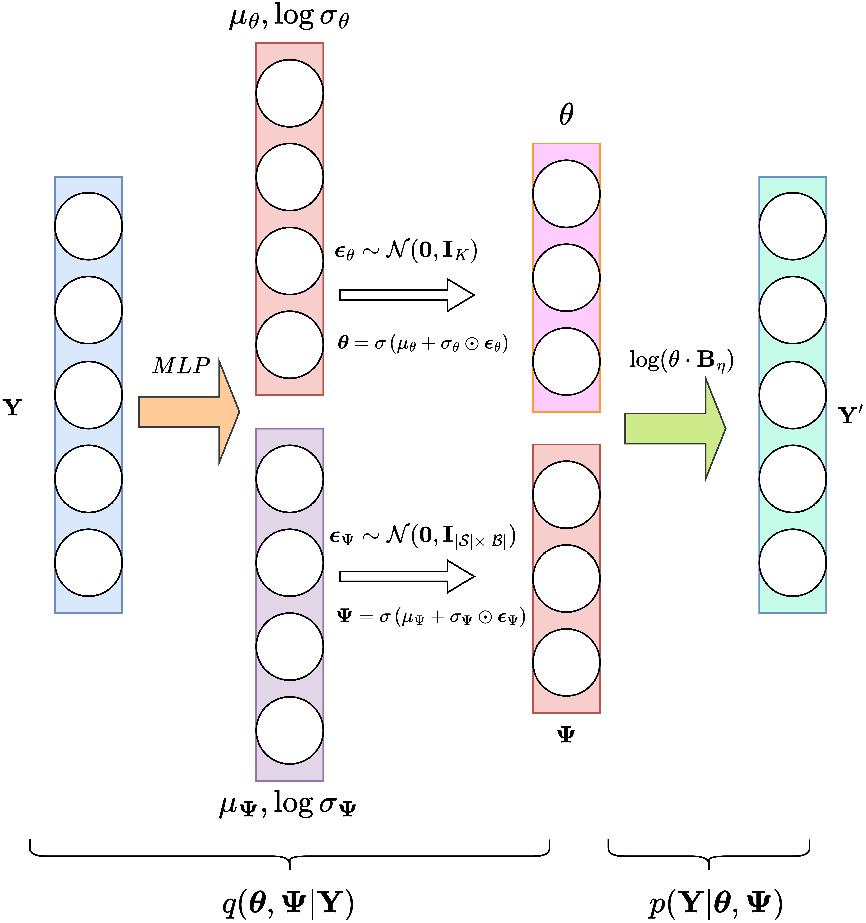
Network structure of inference model *q*(***θ*, Ψ**|**Y**) and generative model *p*(**Y**|***θ*, Ψ**).

Neretva was also benchmarked against the following KIR genotypers: Geny, Locityper, T1K, kir-mapper and PING on 40 Human Pangenome Reference Consortium (HPRC) samples [28]. The KIR ground truth calls for the HPRC samples were obtained from the HPRC assemblies [28] via the BAKIR annotation tool [11].

We evaluated the performance of each tool and gathered both sample-level and allele-level metrics consisting of the number of correct calls, accuracy, precision, recall and *F*_1_ score. On sample level, a call is considered as correct if both (1) allele calls are concordant with ground truth annotation^5^, and (2) copy number is correct. Neretva always produces a unique solution; for tools that produce multiple solutions for a sample, we mark a call as correct if any of the produced solutions is concordant. The accuracy is defined as the number of true positives divided by the number of samples, while the precision is defined as the number of true positives divided by the number of true and false positives. The recall is defined as number of true positives divided by the number of true positives and false negatives.

### Pharmacogene performance

As shown in Table 1 (and Supplementary Table 3), Neretva was able to consistently perfectly genotype *CYP2C19*, and matched the state of the art, such as Aldy, Astrolabe, StellarPGX and PyPGx. Same trends were observed for *CYP2C8*, and *CYP2C9* as well (see Supplementary Table 3). As for *CYP2D6*, Neretva achieved both sample-level and allele-level *F*_1_ of 0.99, putting it on par with Aldy (the second-best tool) and just one call below Cyrius, which achieved perfect accuracy. After further investigation, we observed that our model miscalled *2/*36+*10 as *2/*36+*2 due to the misidentification of a single variant distinguishing *2 and *10. While Aldy was able to solve this case, it failed on another instance with an increased copy number that Neretva was able to handle successfully.

**Table 1.**
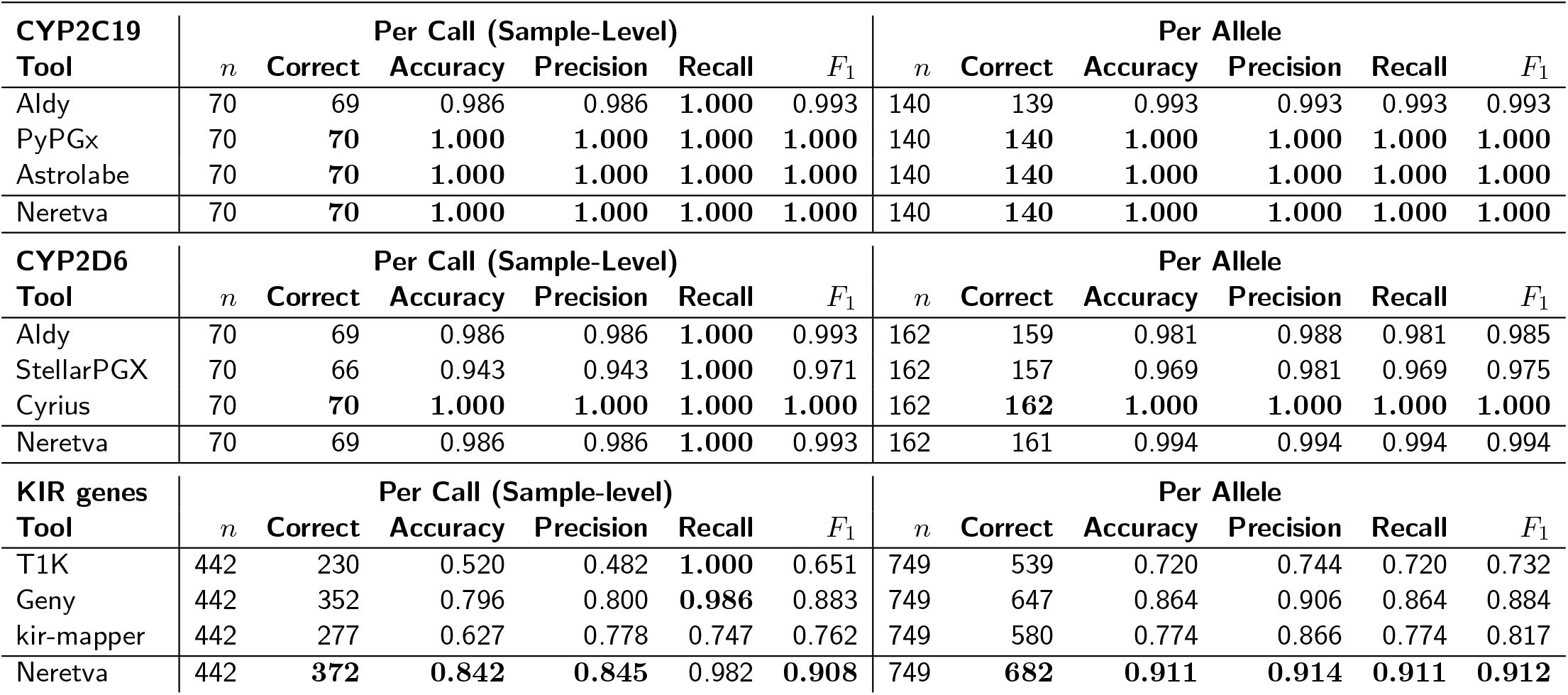
Performance comparison between Neretva and other tools on pharmacogenes and KIR genes. In addition to Neretva, only top three tools are shown here (other tools are shown in Supplementary Table 3). The best results are highlighted in bold.

### KIR gene family performance

As shown in Table 1 (and Supplementary Table 3), Neretva offers improved performance across the evaluated KIR samples in nearly all categories. At the sample level, Neretva achieved the highest accuracy (0.842) and *F*_1_ (0.908), substantially outperforming Geny, the second-best method, which reached a sample accuracy of 0.796. kir-mapper demonstrated competitive precision (0.778) but a noticeably lower recall (0.747), indicating reduced sensitivity. In contrast, T1K achieved lower accuracy on this dataset, with a sample accuracy of 0.520.

At the allele level, Neretva correctly identified 682 out of 749 alleles, achieving an *F*_1_ score of 0.912, thus improving on Geny (allele-level *F*_1_ score: 0.884) and T1K (allele-level *F*_1_ score: 0.732). Per-gene results are available in Supplementary Table These results suggest that Neretva more effectively handles the high sequence similarity and read-mapping ambiguity characteristic of the KIR region, thereby reducing the false positive rates observed in other tools.

Note that Locityper’s performance is significantly lower than originally expected (Supplementary Table 3). We have reached out to the authors for further clarification and guidance. We also note that our evaluation of all tools differs slightly from the results published in Geny [50]. The reason for that is improved handling of the false negatives in Table 1 (in some instances, false negatives were systematically ignored for all tools in Geny’s evaluation script).

### Ablations and computational efficiency

To show the importance of each modelling feature, we performed a set of ablation experiments to demonstrate the impact of the Jensen-Shannon divergence term, entropy penalty, and bias modelling on KIR genes, where all terms are required for optimal performance. The Supplementary Table 1 shows that the *F*_1_ score is highest when all modelling features are enabled.

We also show that Neretva’s approach is more practical than ILP modelling on the most complex dataset (KIR). For this, we devised an ILP model analogous to Neretva and used the state-of-the-art Gurobi solver to solve it (Supplementary Table 2). While Neretva spent less than 8 minutes per model, no ILP was able to converge to the optimal solution in 30 minutes (time limit set to 4× of that of Neretva). The solutions that were obtained were all strictly worse than Neretva, as shown in Supplementary Table 2.

In general, Neretva inference takes no more than a few minutes to complete on pharmacogenes. The more substantial improvements occur on KIR genes; while other tools might take an hour to complete, Neretva typically takes less than 20 minutes. The inference itself takes no more than a few minutes (8 minutes in the worst case); the rest is spent on alignment and data parsing. All experiments were done on an Intel Xeon 8260 machine with 1 TB of RAM and an NVidia Tesla V100 GPU with 32 GB of VRAM.

## Discussion

In this paper, we have presented Neretva, a unified variational inference framework for genotyping highly polymorphic gene families. A key innovation of Neretva is its reformulation of the genotyping problem from combinatorial optimization to variational inference, enabling scalable gradient-based optimization that naturally accommodates uncertainty while maintaining biological interpretability. Unlike ILP approaches that guarantee optimality only under rigid model assumptions and lack scalability, we have shown through extensive experiments across major complex gene families that Neretva’s probabilistic framework naturally handles uncertainty and can scale to large gene families with complex features.

Several limitations of the current version of Neretva warrant discussion and further exploration. First, the current model treats variant observations as independent counts at each position and does not consider read-level linkage information. We hope to extend the model to incorporate read-level information. Secondly, support for various KIR fusions involving one or more genes, in addition to certain complex structural variations within the *CYP2D6* locus, still requires improvements in copy number modelling and a more extensive evaluation dataset (while Neretva can call gene copy numbers and some fusions, the accuracy is currently sensitive to sequencing noise).

A few promising directions merit future investigation. The first is the support for long-read sequencing platforms such as Oxford Nanopore. Moreover, the generality of our framework suggests a potential inclusion of other clinically important gene families, such as HLA and T-cell receptor genes. Another promising direction is using pangenome graph references to obtain alignments and model base counts. Recent advances in graph neural networks, including variational graph auto-encoders [23], also provide principled approaches for learning representations over such structures, and we aim to explore their integration to better reason about overlapping haplotypes and complex structural variation. Finally, our reliance on open-source libraries enables us to explore the potential integration with privacy-preserving technologies that provide support for secure tensor algebra in the near future.

Our work demonstrates that Neretva achieves competitive or improved accuracy compared to existing tools, especially on more complex gene families such as KIR, and that its variational inference framework provides a principled and extensible foundation for genotyping complex genomic loci. We hope that our work, however modest, contributes to improved decisions in precision medicine.

## Funding

This work was supported by NSERC Discovery RGPIN-2019-04973 and RGPIN-2025-04073, Canada Research Chairs, Canada Foundation for Innovation and BC Knowledge Development Fund grants.

## Conflicts of interest

None declared.

## Data availability

Neretva is open source and is freely available at https://github.com/0xTCG/neretva. This repository contains all tool outputs, as well as a detailed description of the results and data sources. Data sources are listed in Appendix.

## Author contributions

Q.Z. conceived the study and designed and implemented the framework. I.N. oversaw the study. S.P.A. conducted the experiments. All authors jointly wrote and reviewed the manuscript.

## Footnotes

1 For example, Aldy was shown to consistently produce good results across many different HTS technologies, often achieving up to 99% accuracy

2 By using -x sr --secondary=yes -c --dual=no parameters.

3 Other pharmacogenes were not considered to avoid clutter, and also because their complexity does not surpass the complexity of the families evaluated here [15, 14]

4 While GeT-RM project describes 137 samples, only 70 of them are publicly available in the evaluated WGS cohort.

5 Concordance means that the allele call matches the ground truth at functional level; in case of pharmacogenes, that implies the major star-allele match (e.g., *2A is regarded as *2), while for KIR genes, the match of the first 3 digits within the allele name (e.g., *0030201 is regarded as *003).

## Notes

https://github.com/0xTCG/neretva

### Competing Interest Statement

The authors have declared no competing interest.

### Summary of Updates

- Streamlined text - New results and ablations

