## Appendix for "Neretva: Neural Variational Inference for Allele-level Genotyping of Highly Polymorphic Genes"

Qinghui Zhou, Seyed Pouria Ahmadi and and Ibrahim Numanagić

### 1 Deriving the Evidence Lower Bound

Let us derive the evidence lower bound (ELBO) for our variational inference objective. From the log-marginal likelihood:

$$\log p(\mathbf{Y}) = \log \int_\theta \int_\Psi p(\mathbf{Y}|\boldsymbol{\theta}, \boldsymbol{\Psi})p(\boldsymbol{\theta})p(\boldsymbol{\Psi})\, d\boldsymbol{\theta}\, d\boldsymbol{\Psi},$$

we introduce the variational distribution $q(\boldsymbol{\theta}, \boldsymbol{\Psi}|\mathbf{Y})$ and apply Jensen's inequality:

$$\begin{aligned}
\log p(\mathbf{Y}) &= \log \int_\theta \int_\Psi p(\mathbf{Y}|\boldsymbol{\theta}, \boldsymbol{\Psi})p(\boldsymbol{\theta})p(\boldsymbol{\Psi}) \frac{q(\boldsymbol{\theta}, \boldsymbol{\Psi}|\mathbf{Y})}{q(\boldsymbol{\theta}, \boldsymbol{\Psi}|\mathbf{Y})} d\boldsymbol{\theta}\, d\boldsymbol{\Psi} \\
&= \log \mathbb{E}_{q(\boldsymbol{\theta}, \boldsymbol{\Psi}|\mathbf{Y})} \left[ \frac{p(\mathbf{Y}|\boldsymbol{\theta}, \boldsymbol{\Psi})p(\boldsymbol{\theta})p(\boldsymbol{\Psi})}{q(\boldsymbol{\theta}, \boldsymbol{\Psi}|\mathbf{Y})} \right] \\
&\geq \mathbb{E}_{q(\boldsymbol{\theta}, \boldsymbol{\Psi}|\mathbf{Y})} \left[ \log \frac{p(\mathbf{Y}|\boldsymbol{\theta}, \boldsymbol{\Psi})p(\boldsymbol{\theta})p(\boldsymbol{\Psi})}{q(\boldsymbol{\theta}, \boldsymbol{\Psi}|\mathbf{Y})} \right].
\end{aligned}$$

After expanding the logarithm, we obtain:

$$\begin{aligned}
\log p(\mathbf{Y}) \geq\ & \mathbb{E}_q[\log p(\mathbf{Y}|\boldsymbol{\theta}, \boldsymbol{\Psi})] + \mathbb{E}_q[\log p(\boldsymbol{\theta})] \\
&+ \mathbb{E}_q[\log p(\boldsymbol{\Psi})] - \mathbb{E}_q[\log q(\boldsymbol{\theta}, \boldsymbol{\Psi}|\mathbf{Y})].
\end{aligned}$$

Under the mean-field approximation as specified in (2), we get:

$$\mathbb{E}_q[\log q(\boldsymbol{\theta}, \boldsymbol{\Psi}|\mathbf{Y})] = \mathbb{E}_q[\log q(\boldsymbol{\theta}|\mathbf{Y})] + \sum_{s \in \mathcal{S}} \mathbb{E}_q[\log q(\boldsymbol{\psi}_s|\mathbf{Y})]$$

In a similar vein, for the prior $p(\boldsymbol{\Psi}) = \prod_{s \in \mathcal{S}} p(\boldsymbol{\psi}_s)$, we get:

$$\mathbb{E}_q[\log p(\boldsymbol{\Psi})] = \sum_{s \in \mathcal{S}} \mathbb{E}_q[\log p(\boldsymbol{\psi}_s)].$$

Finally, by rearranging the terms, we obtain the final ELBO:

$$\begin{aligned}
\mathcal{L} = \mathbb{E}_q[\log p(\mathbf{Y}|\boldsymbol{\theta}, \boldsymbol{\Psi})] &- D_{\mathsf{KL}}[q(\boldsymbol{\theta}|\mathbf{Y}) \| p(\boldsymbol{\theta})] \\
&- \sum_{s \in \mathcal{S}} D_{\mathsf{KL}}[q(\boldsymbol{\psi}_s|\mathbf{Y}) \| p(\boldsymbol{\psi}_s)].
\end{aligned}$$

#### 1.1 Reparameterization and Inference

To optimize this objective, we parameterize the variational posterior using an encoder network with parameters $\phi$:

$$(\boldsymbol{\mu}_\theta, \log \boldsymbol{\sigma}^2_\theta, \boldsymbol{\mu}_\Psi, \log \boldsymbol{\sigma}^2_\Psi) = \mathsf{EncoderNeuralNet}_\phi(\mathbf{Y}).$$

To enable gradient-based optimization through the stochastic layers, apply the reparameterization trick (Kingma and Welling, 2013). First, sample noise from standard Gaussians:

$$\begin{aligned}
\boldsymbol{\epsilon}_\theta &\sim \mathcal{N}(\mathbf{0}, \mathbf{I}_K); \\
\boldsymbol{\epsilon}_\Psi &\sim \mathcal{N}(\mathbf{0}, \mathbf{I}_{|\mathcal{S}| \times |\mathcal{B}|}).
\end{aligned}$$

Then, transform these to obtain samples from the variational posterior:

$$\begin{aligned}
\boldsymbol{\theta} &= \sigma\left(\mu_\theta + \boldsymbol{\sigma_\theta} \odot \boldsymbol{\epsilon}_\theta\right); \\
\boldsymbol{\Psi} &= \sigma\left(\mu_\Psi + \boldsymbol{\sigma_\Psi} \odot \boldsymbol{\epsilon}_\Psi\right).
\end{aligned}$$

This yields the variational posteriors:

$$\begin{aligned}
q(\boldsymbol{\theta}|\mathbf{Y}) &= \mathcal{LN}(\mu_\theta, \mathsf{diag}(\sigma^2_\theta)) \\
q(\boldsymbol{\Psi}|\mathbf{Y}) &= \mathcal{LN}(\mu_\Psi, \mathsf{diag}(\sigma^2_\Psi)),
\end{aligned}$$

where $\sigma(\cdot)$ denotes the softmax function. This reparametrization allows us to optimize the ELBO using standard backpropagation with the SGVB estimator (Kingma and Welling, 2013) via a single Monte Carlo sample.

#### 2 Model Parameters

- Copy number model: 500 optimizer iterations for pharmacogenes and 1000 iterations for KIR. KIR requires more copy number iterations due to its larger search space spanning 17 genes with independent copy numbers.
- Copy number model: $\delta = 0.5$ for pharmacogenes and $\delta = 0.3$ for KIR. $\delta$ is lower for KIR to increase robustness to the greater coverage variability arising from cross-gene mappings common in KIR.
- Bias modelling was only used for KIR genes due to ambiguous read mappings. It was not used for pharmacogenes which do not need it.
- ELBO model: For KIR, $\beta = 600,000$ and $\gamma = 100$. Complex gene families, such as KIR and *CYP2D6*, define more minor variants than core variants. As reconstruction loss is dominated by minor variant signal, a larger $\beta$ helps regularize minor variants and emphasize the core variants. For *CYP2D6*, used a warm up schedule for $\beta$ between $0$ and $600,000$. For *CYP2D6*, a warm-up schedule for $\beta$ was found to improve accuracy empirically; for KIR, a high $\beta$ performed best so no warmup schedule was set up for KIR (warmup is still possible but at the price of slower runtime). For other (simpler) pharmacogenes, we used a warm up schedule for $\beta$ between $0$ and $0.6$, and $\gamma = 100$.
- ELBO model: 3,000 optimizer iterations on both pharmacogenes and KIR.
- Learning rate: 0.005 for allele parameters, and 0.00001 for base parameters.
- Random seeds: 42 for KIR, and multi-seeds (42, 123, 456) for pharmacogenes with best loss selection. The smaller instance of the pharmacogene model permits multiple seed selection at negligible additional cost, while KIR's larger model makes multi-seed runs costly.

#### 3 Software Versions

We used StellarPGX (v1.2.7), Aldy (v4.8.2), Cyrius (v1.1.1), Stargazer (v1.0.9), Astrolabe (v0.8.7.2) and PyPGx (v0.25.0). For KIR genes, we used Geny (v1.0), Locityper (v1.2.3), T1K (v1.0.9), kir-mapper (v1.01) and PING (commit b4b6957) on 40 Human Pangenome Reference Consortium (HPRC) samples (Liao et al., 2023).

#### 4 Data Availability

Neretva is open source and is freely available at https://github.com/0xTCG/neretva. This repository contains all tool outputs, as well as a detailed description of the results and data sources. The complete sequencing data is available at https://github.com/Illumina/Polaris/wiki/HiSeqX-PGx-Cohort and https://humanpangenome.org/data/.

### 5 Ablation Studies

#### 5.1 Modelling Features

Supplementary Table 1 shows the results of the Neretva model with a single feature turned off, in addition to the full model (bottom row).

**Supplemental Table 1:** Ablation study on KIR genes (HPRC dataset). Each row removes one component from the full Neretva model (Equation 4). $\beta$=0: Jensen-Shannon divergence term removed; $\gamma$=0: entropy penalty removed; $\mathbf{B}_\eta$=$\mathbf{B}$: bias modeling disabled. The best results are highlighted in bold.

| Model | Per Call (Sample-Level) | | | | | | Per Allele | | | | | |
|---|---|---|---|---|---|---|---|---|---|---|---|---|
| Variant | $n$ | Correct | Accuracy | Precision | Recall | $F_1$ | $n$ | Correct | Accuracy | Precision | Recall | $F_1$ |
| $\beta$=0 | 442 | 328 | 0.742 | 0.744 | 0.979 | 0.845 | 749 | 616 | 0.822 | 0.834 | 0.822 | 0.828 |
| $\gamma$=0 | 442 | 371 | 0.839 | 0.841 | **0.984** | 0.907 | 749 | **684** | **0.913** | 0.908 | **0.913** | 0.911 |
| $\mathbf{B}_\eta$=$\mathbf{B}$ | 442 | 365 | 0.826 | 0.851 | 0.955 | 0.900 | 749 | 672 | 0.897 | **0.916** | 0.897 | 0.906 |
| Neretva | 442 | **372** | **0.842** | **0.845** | 0.982 | **0.908** | 749 | 682 | 0.911 | 0.914 | 0.911 | **0.912** |

#### 5.2 Integer Linear Programming Formulation

To evaluate whether the continuous relaxation provided by the AEVB is necessary, we show an analogous integer linear program (ILP) below. The results of this formulation were shown in Supplementary Table 2.

**Supplemental Table 2:** Comparison of Neretva (AEVB) and an analogous ILP formulation (see the last section in Appendix) on KIR genes (HPRC dataset). The ILP solver was given a 1800 s timeout per sample; "Crashed" denotes samples where the solver timed out with an optimality gap exceeding 80%. The best results are highlighted in bold.

| | Per Call (Sample-Level) | | | | | | Per Allele | | | | | | |
|---|---|---|---|---|---|---|---|---|---|---|---|---|---|
| **Method** | $n$ | Correct | Accuracy | Precision | Recall | $F_1$ | $n$ | Correct | Accuracy | Precision | Recall | $F_1$ | Crashed |
| ILP | 442 | 248 | 0.561 | 0.598 | 0.879 | 0.712 | 749 | 558 | 0.745 | 0.772 | 0.745 | 0.758 | 11/40 |
| Neretva | 442 | **372** | **0.842** | **0.845** | **0.982** | **0.908** | 749 | **682** | **0.911** | **0.914** | **0.911** | **0.912** | - |

The ILP contains a median of 228,729 binary variables and 732,250 constraints across the 40 HPRC samples, with the largest instance reaching over 541,000 binary variables and 1.7 million constraints. We solved each instance using Gurobi 12.0 with a time limit of 1,800 seconds, approximately four times the maximum AEVB runtime of 447 seconds (mean: 190 seconds). Despite this generous time budget, all 40 instances hit the time limit without reaching optimality, with a median optimality gap of 43.9% and 20% of samples exceeding a 95% gap. The best feasible solutions achieved an allele-level F1 score of 0.758, compared to 0.912 for the AEVB. These results demonstrate that the discrete formulation is both computationally intractable and produces inferior genotyping accuracy, validating the necessity of the AEVB framework.

##### Index Sets

- $\mathcal{A}$: set of candidate alleles, indexed by $a$
- $\mathcal{P}_a$: set of generatable positions for allele $a$, indexed by $p$
- $\mathcal{B} = \{$A, C, G, T, N, P$\}$: set of bases, indexed by $k$
- $\mathcal{M}$: set of mutation indices, indexed by $m$

##### Data

- $y_m \in \mathbb{N}$: observed read count at mutation index $m$
- $d$: expected per-copy sequencing depth (i.e., one copy of an allele contributes $d$ reads at each of its positions)
- $\sigma : (\mathcal{A}, \mathcal{P}, \mathcal{B}) \to \mathcal{M}$: mapping from (allele, position, base) triples to mutation indices

- $\gamma_{a,p,k} \in \mathbb{R}$: cost of allele $a$ selecting base $k$ at position $p$. Selecting a different base from database expectation incurs higher cost.

##### Decision Variables

| | |
|---|---|
| $c_a \in \{0,1,2,3,4\}$ | copy number of allele $a$ |
| $b_{a,p,k} \in \{0,1\}$ | 1 if allele $a$ selects base $k$ at position $p$ |
| $z_{a,p,k} \in [0,4]$ | linearized product: $z_{a,p,k} = c_a \cdot b_{a,p,k}$ |
| $e_m^+,\ e_m^- \geq 0$ | positive/negative reconstruction error at mutation $m$ |

##### Constraints

*Base selection.* Each allele selects exactly one base per generatable position:

$$\sum_{k \in \mathcal{B}} b_{a,p,k} = 1 \qquad \forall\, a \in \mathcal{A},\ p \in \mathcal{P}_a$$

*Linearization of $z_{a,p,k} = c_a \cdot b_{a,p,k}$.* Standard McCormick envelope (where $C_{\max} = 4$ is the maximum copy number):

$$\begin{aligned} z_{a,p,k} &\leq C_{\max} \cdot b_{a,p,k} \\ z_{a,p,k} &\leq c_a \\ z_{a,p,k} &\geq c_a - C_{\max}(1 - b_{a,p,k}) \end{aligned}$$

*Reconstruction.* The observed count at each mutation index equals the sum of contributions from all alleles plus error terms:

$$d \sum_{\substack{(a,p,k) \\ \sigma(a,p,k)=m}} z_{a,p,k} \;+\; e_m^- \;-\; e_m^+ \;=\; y_m \qquad \forall\, m \in \mathcal{M}$$

##### Objective

$$\min \quad \sum_{m \in \mathcal{M}} \left(e_m^+ + e_m^-\right) \;+\; \lambda \sum_{a \in \mathcal{A}} \sum_{p \in \mathcal{P}_a} \sum_{k \in \mathcal{B}} \gamma_{a,p,k} \cdot b_{a,p,k}$$

where $\lambda > 0$ controls the trade-off between reconstruction fidelity and adherence to the database prior.

### 6 Supplementary Results

**Supplemental Table 3:** Complete performance comparison between Neretva and other tools on pharmacogenes and KIR genes. The best results are highlighted in bold.

| **CYP2C8** | | | **Per Call (Sample-Level)** | | | | | | **Per Allele** | | | |
|---|---|---|---|---|---|---|---|---|---|---|---|---|
| **Tool** | $n$ | **Correct** | **Accuracy** | **Precision** | **Recall** | $F_1$ | $n$ | **Correct** | **Accuracy** | **Precision** | **Recall** | $F_1$ |
| Aldy | 70 | **70** | **1.000** | **1.000** | **1.000** | **1.000** | 140 | **140** | **1.000** | **1.000** | **1.000** | **1.000** |
| Stargazer | **70** | 68 | 0.971 | 0.971 | **1.000** | 0.986 | 140 | 138 | 0.986 | 0.986 | 0.986 | 0.986 |
| PyPGx | 70 | **70** | **1.000** | **1.000** | **1.000** | **1.000** | 140 | **140** | **1.000** | **1.000** | **1.000** | **1.000** |
| Astrolabe | 70 | **70** | **1.000** | **1.000** | **1.000** | **1.000** | 140 | **140** | **1.000** | **1.000** | **1.000** | **1.000** |
| Neretva | 70 | **70** | **1.000** | **1.000** | **1.000** | **1.000** | 140 | **140** | **1.000** | **1.000** | **1.000** | **1.000** |

| **CYP2C9** | | | **Per Call (Sample-Level)** | | | | | | **Per Allele** | | | |
|---|---|---|---|---|---|---|---|---|---|---|---|---|
| **Tool** | $n$ | **Correct** | **Accuracy** | **Precision** | **Recall** | $F_1$ | $n$ | **Correct** | **Accuracy** | **Precision** | **Recall** | $F_1$ |
| Aldy | 70 | **70** | **1.000** | **1.000** | **1.000** | **1.000** | 140 | **140** | **1.000** | **1.000** | **1.000** | **1.000** |
| StellarPGx | 70 | **70** | **1.000** | **1.000** | **1.000** | **1.000** | 140 | **140** | **1.000** | **1.000** | **1.000** | **1.000** |
| Stargazer | 70 | 68 | 0.971 | 0.971 | **1.000** | 0.986 | 140 | 138 | 0.986 | 0.986 | 0.986 | 0.986 |
| PyPGx | 70 | 68 | 0.971 | 0.971 | **1.000** | 0.986 | 140 | 138 | 0.986 | 0.986 | 0.986 | 0.986 |
| Astrolabe | 70 | 67 | 0.957 | 0.957 | **1.000** | 0.978 | 140 | 137 | 0.979 | 0.979 | 0.979 | 0.979 |
| Neretva | 70 | **70** | **1.000** | **1.000** | **1.000** | **1.000** | 140 | **140** | **1.000** | **1.000** | **1.000** | **1.000** |

| **CYP2C19** | | | **Per Call (Sample-Level)** | | | | | | **Per Allele** | | | |
|---|---|---|---|---|---|---|---|---|---|---|---|---|
| **Tool** | $n$ | **Correct** | **Accuracy** | **Precision** | **Recall** | $F_1$ | $n$ | **Correct** | **Accuracy** | **Precision** | **Recall** | $F_1$ |
| Aldy | 70 | 69 | 0.986 | 0.986 | **1.000** | 0.993 | 140 | 139 | 0.993 | 0.993 | 0.993 | 0.993 |
| StellarPGx | 70 | 69 | 0.986 | 0.986 | **1.000** | 0.993 | 140 | 139 | 0.993 | 0.993 | 0.993 | 0.993 |
| Stargazer | 70 | 69 | 0.986 | 0.986 | **1.000** | 0.993 | 140 | 139 | 0.993 | 0.993 | 0.993 | 0.993 |
| PyPGx | 70 | **70** | **1.000** | **1.000** | **1.000** | **1.000** | 140 | **140** | **1.000** | **1.000** | **1.000** | **1.000** |
| Astrolabe | 70 | **70** | **1.000** | **1.000** | **1.000** | **1.000** | 140 | **140** | **1.000** | **1.000** | **1.000** | **1.000** |
| Neretva | 70 | **70** | **1.000** | **1.000** | **1.000** | **1.000** | 140 | **140** | **1.000** | **1.000** | **1.000** | **1.000** |

| **CYP2D6** | | | **Per Call (Sample-Level)** | | | | | | **Per Allele** | | | |
|---|---|---|---|---|---|---|---|---|---|---|---|---|
| **Tool** | $n$ | **Correct** | **Accuracy** | **Precision** | **Recall** | $F_1$ | $n$ | **Correct** | **Accuracy** | **Precision** | **Recall** | $F_1$ |
| Aldy | 70 | 69 | 0.986 | 0.986 | **1.000** | 0.993 | 162 | 159 | 0.981 | 0.988 | 0.981 | 0.985 |
| StellarPGX | 70 | 66 | 0.943 | 0.943 | **1.000** | 0.971 | 162 | 157 | 0.969 | 0.981 | 0.969 | 0.975 |
| Cyrius | 70 | **70** | **1.000** | **1.000** | **1.000** | **1.000** | 162 | **162** | **1.000** | **1.000** | **1.000** | **1.000** |
| Stargazer | 70 | 40 | 0.571 | 0.571 | **1.000** | 0.727 | 162 | 123 | 0.759 | 0.879 | 0.759 | 0.815 |
| PyPGx | 70 | 37 | 0.529 | 0.529 | **1.000** | 0.692 | 162 | 122 | 0.753 | 0.871 | 0.753 | 0.808 |
| Astrolabe | 70 | 36 | 0.514 | 0.514 | **1.000** | 0.679 | 162 | 120 | 0.741 | 0.857 | 0.741 | 0.795 |
| Neretva | 70 | 69 | 0.986 | 0.986 | **1.000** | 0.993 | 162 | 161 | 0.994 | 0.994 | 0.994 | 0.994 |

| **KIR genes** | | | **Per Call (Sample-level)** | | | | | | **Per Allele** | | | |
|---|---|---|---|---|---|---|---|---|---|---|---|---|
| **Tool** | $n$ | **Correct** | **Accuracy** | **Precision** | **Recall** | $F_1$ | $n$ | **Correct** | **Accuracy** | **Precision** | **Recall** | $F_1$ |
| PING | 442 | 229 | 0.518 | 0.577 | 0.640 | 0.607 | 749 | 464 | 0.619 | 0.699 | 0.619 | 0.657 |
| T1K | 442 | 230 | 0.520 | 0.482 | **1.000** | 0.651 | 749 | 539 | 0.720 | 0.744 | 0.720 | 0.732 |
| Geny | 442 | 352 | 0.796 | 0.800 | **0.986** | 0.883 | 749 | 647 | 0.864 | 0.906 | 0.864 | 0.884 |
| kir-mapper | 442 | 277 | 0.627 | 0.778 | 0.747 | 0.762 | 749 | 580 | 0.774 | 0.866 | 0.774 | 0.817 |
| Locityper | 442 | 59 | 0.133 | 0.107 | 0.431 | 0.172 | 749 | 343 | 0.458 | 0.311 | 0.458 | 0.371 |
| Neretva | 442 | **372** | **0.842** | **0.845** | 0.982 | **0.908** | 749 | **682** | **0.911** | **0.914** | **0.911** | **0.912** |

**Supplemental Table 4:** Per-gene performance comparison on KIR genes (HPRC dataset). The best results per gene are highlighted in bold.

#### (a) Per Call (Sample-Level)

| | | PING | | | | T1K | | | | Geny | | | | kir-mapper | | | | Locityper | | | | Neretva | | | |
|---|---|---|---|---|---|---|---|---|---|---|---|---|---|---|---|---|---|---|---|---|---|---|---|---|---|
| **Gene** | **Total** | **Correct** | **Precision** | **Recall** | $F_1$ | **Correct** | **Precision** | **Recall** | $F_1$ | **Correct** | **Precision** | **Recall** | $F_1$ | **Correct** | **Precision** | **Recall** | $F_1$ | **Correct** | **Precision** | **Recall** | $F_1$ | **Correct** | **Precision** | **Recall** | $F_1$ |
| *KIR2DL1* | 39 | 19 | **90.5%** | 51.4% | 0.66 | 15 | 38.5% | **100.0%** | 0.56 | 28 | 71.8% | **100.0%** | 0.84 | 16 | 50.0% | 69.6% | 0.58 | 3 | 8.3% | 42.9% | 0.14 | **32** | 82.1% | **100.0%** | **0.90** |
| *KIR2DL2* | 17 | 3 | 8.6% | 50.0% | 0.15 | 8 | 47.1% | **100.0%** | 0.64 | **16** | **94.1%** | **100.0%** | **0.97** | 4 | 57.1% | 28.6% | 0.38 | 2 | 5.4% | **100.0%** | 0.10 | 15 | 93.8% | 93.8% | 0.94 |
| *KIR2DL3* | 37 | 19 | 54.3% | 82.6% | 0.66 | 21 | 55.3% | **100.0%** | 0.71 | 31 | 83.8% | **100.0%** | 0.91 | 18 | 81.8% | 54.5% | 0.65 | 19 | 47.5% | **100.0%** | 0.64 | **35** | **94.6%** | **100.0%** | **0.97** |
| *KIR2DL4* | 39 | 29 | **87.9%** | 82.9% | 0.85 | 21 | 53.8% | **100.0%** | 0.70 | **34** | 87.2% | **100.0%** | **0.93** | 31 | 86.1% | 91.2% | 0.89 | 3 | 15.8% | 12.5% | 0.14 | **34** | 87.2% | **100.0%** | **0.93** |
| *KIR2DL5A* | 7 | 5 | 33.3% | 71.4% | 0.45 | 2 | 15.4% | **100.0%** | 0.27 | **7** | **70.0%** | **100.0%** | **0.82** | 5 | 45.5% | **100.0%** | 0.62 | 0 | 0.0% | 0.0% | — | 6 | 66.7% | **100.0%** | 0.80 |
| *KIR2DL5B* | 12 | **9** | 60.0% | 81.8% | 0.69 | 8 | 22.2% | **100.0%** | 0.36 | 8 | **80.0%** | 80.0% | **0.80** | 6 | 40.0% | 85.7% | 0.55 | 0 | 0.0% | 0.0% | — | 8 | 66.7% | 88.9% | 0.76 |
| *KIR2DP1* | 39 | 19 | **95.0%** | 50.0% | 0.66 | 18 | 46.2% | **100.0%** | 0.63 | 30 | 78.9% | 96.8% | 0.87 | 25 | 78.1% | 78.1% | 0.78 | 7 | 18.4% | 77.8% | 0.30 | **34** | 89.5% | 97.1% | **0.93** |
| *KIR2DS1* | 12 | 0 | 0.0% | 0.0% | — | 8 | 66.7% | **100.0%** | 0.80 | **12** | **100.0%** | **100.0%** | **1.00** | 8 | 80.0% | 80.0% | 0.80 | 2 | 5.0% | **100.0%** | 0.10 | 10 | 90.9% | 90.9% | 0.91 |
| *KIR2DS2* | 16 | 13 | 92.9% | 86.7% | 0.90 | 12 | 75.0% | **100.0%** | 0.86 | **15** | 93.8% | **100.0%** | **0.97** | 3 | 75.0% | 20.0% | 0.32 | 0 | 0.0% | — | — | **15** | **100.0%** | 93.8% | **0.97** |
| *KIR2DS3* | 7 | 6 | 33.3% | **100.0%** | 0.50 | 4 | 40.0% | **100.0%** | 0.57 | 6 | **100.0%** | 85.7% | **0.92** | 1 | **100.0%** | 14.3% | 0.25 | 0 | 0.0% | — | — | **7** | 77.8% | **100.0%** | 0.88 |
| *KIR2DS4* | 38 | 30 | 88.2% | 88.2% | 0.88 | 24 | 63.2% | **100.0%** | 0.77 | 32 | 84.2% | **100.0%** | 0.91 | 28 | 90.3% | 80.0% | 0.85 | 1 | 6.2% | 4.0% | 0.05 | **36** | **94.7%** | **100.0%** | **0.97** |
| *KIR2DS5* | 12 | 8 | 44.4% | 88.9% | 0.59 | 8 | 66.7% | **100.0%** | 0.80 | **11** | **100.0%** | 91.7% | **0.96** | 2 | 50.0% | 20.0% | 0.29 | 0 | 0.0% | 0.0% | — | 10 | **100.0%** | 83.3% | 0.91 |
| *KIR3DL1* | 38 | 27 | 73.0% | 93.1% | 0.82 | 31 | 81.6% | **100.0%** | 0.90 | **34** | 89.5% | **100.0%** | **0.94** | 27 | **96.4%** | 73.0% | 0.83 | 0 | 0.0% | 0.0% | — | 32 | 84.2% | **100.0%** | 0.91 |
| *KIR3DL2* | 40 | 0 | 0.0% | 0.0% | — | 22 | 55.0% | **100.0%** | 0.71 | 32 | 80.0% | **100.0%** | 0.89 | **36** | **97.3%** | 92.3% | **0.95** | 3 | 11.5% | 17.6% | 0.14 | 32 | 80.0% | **100.0%** | 0.89 |
| *KIR3DL3* | 40 | 24 | 72.7% | 77.4% | 0.75 | 15 | 37.5% | **100.0%** | 0.55 | 22 | 55.0% | **100.0%** | 0.71 | **31** | **79.5%** | 96.9% | 0.87 | 8 | 20.5% | 88.9% | 0.33 | **31** | 77.5% | **100.0%** | **0.87** |
| *KIR3DP1* | 39 | 13 | 54.2% | 46.4% | 0.50 | 7 | 17.5% | **100.0%** | 0.30 | 25 | 64.1% | **100.0%** | 0.78 | **29** | **78.4%** | 93.5% | **0.85** | 9 | 23.1% | 90.0% | 0.37 | 25 | 64.1% | **100.0%** | 0.78 |
| *KIR3DS1* | 10 | 5 | 13.5% | **100.0%** | 0.24 | 6 | 60.0% | **100.0%** | 0.75 | 9 | 90.0% | **100.0%** | 0.95 | 7 | 70.0% | **100.0%** | 0.82 | 2 | 5.0% | **100.0%** | 0.10 | **10** | **100.0%** | **100.0%** | **1.00** |
| All | 442 | 229 | 57.7% | 64.0% | 0.61 | 230 | 48.2% | **100.0%** | 0.65 | 352 | 80.0% | 98.6% | 0.88 | 277 | 77.8% | 74.7% | 0.76 | 59 | 10.7% | 43.1% | 0.17 | **372** | **84.5%** | 98.2% | **0.91** |

#### (b) Per Allele

| | | PING | | | | T1K | | | | Geny | | | | kir-mapper | | | | Locityper | | | | Neretva | | | |
|---|---|---|---|---|---|---|---|---|---|---|---|---|---|---|---|---|---|---|---|---|---|---|---|---|---|
| **Gene** | **Total** | **Correct** | **Precision** | **Recall** | $F_1$ | **Correct** | **Precision** | **Recall** | $F_1$ | **Correct** | **Precision** | **Recall** | $F_1$ | **Correct** | **Precision** | **Recall** | $F_1$ | **Correct** | **Precision** | **Recall** | $F_1$ | **Correct** | **Precision** | **Recall** | $F_1$ |
| *KIR2DL1* | 68 | 37 | **94.9%** | 54.4% | 0.69 | 43 | 75.4% | 63.2% | 0.69 | 56 | 87.5% | 82.4% | 0.85 | 43 | 67.2% | 63.2% | 0.65 | 22 | 30.6% | 32.4% | 0.31 | **65** | 90.3% | **95.6%** | **0.93** |
| *KIR2DL2* | 19 | 14 | 20.9% | 73.7% | 0.33 | 14 | 60.9% | 73.7% | 0.67 | 18 | **94.7%** | 94.7% | **0.95** | 7 | 63.6% | 36.8% | 0.47 | **19** | 25.7% | **100.0%** | 0.41 | 17 | 94.4% | 89.5% | 0.92 |
| *KIR2DL3* | 60 | 50 | 74.6% | 83.3% | 0.79 | 44 | 86.3% | 73.3% | 0.79 | 54 | 94.7% | 90.0% | 0.92 | 40 | 90.9% | 66.7% | 0.77 | 55 | 68.8% | 91.7% | 0.79 | **58** | **100.0%** | **96.7%** | **0.98** |
| *KIR2DL4* | 78 | 62 | 96.9% | 79.5% | 0.87 | 58 | 89.2% | 74.4% | 0.81 | **73** | **100.0%** | **93.6%** | **0.97** | 70 | 97.2% | 89.7% | 0.93 | 20 | 55.6% | 25.6% | 0.35 | **73** | 93.6% | **93.6%** | 0.94 |
| *KIR2DL5A* | 8 | 6 | 33.3% | 75.0% | 0.46 | 5 | 31.2% | 62.5% | 0.42 | **8** | 72.7% | **100.0%** | **0.84** | 6 | 60.0% | 75.0% | 0.67 | 5 | 13.2% | 62.5% | 0.22 | 7 | **77.8%** | 87.5% | 0.82 |
| *KIR2DL5B* | 14 | 11 | 61.1% | 78.6% | 0.69 | **13** | 31.7% | **92.9%** | 0.47 | 10 | **100.0%** | 71.4% | **0.83** | **13** | 61.9% | **92.9%** | 0.74 | 7 | 19.4% | 50.0% | 0.28 | 10 | 76.9% | 71.4% | 0.74 |
| *KIR2DP1* | 69 | 34 | **97.1%** | 49.3% | 0.65 | 48 | 77.4% | 69.6% | 0.73 | 60 | 92.3% | 87.0% | 0.90 | 54 | 85.7% | 78.3% | 0.82 | 40 | 52.6% | 58.0% | 0.55 | **65** | 94.2% | **94.2%** | **0.94** |
| *KIR2DS1* | 14 | 1 | **100.0%** | 7.1% | 0.13 | 11 | 84.6% | 78.6% | 0.81 | **14** | **100.0%** | **100.0%** | **1.00** | 12 | 85.7% | 85.7% | 0.86 | **14** | 17.5% | **100.0%** | 0.30 | 13 | 92.9% | 92.9% | 0.93 |
| *KIR2DS2* | 17 | 13 | 92.9% | 76.5% | 0.84 | 14 | 82.4% | 82.4% | 0.82 | **16** | 94.1% | **94.1%** | 0.94 | 4 | 66.7% | 23.5% | 0.35 | **16** | 20.0% | **94.1%** | 0.33 | **16** | **100.0%** | **94.1%** | **0.97** |
| *KIR2DS3* | 8 | 7 | 35.0% | 87.5% | 0.50 | 7 | 58.3% | 87.5% | 0.70 | 7 | **100.0%** | 87.5% | **0.93** | 2 | **100.0%** | 25.0% | 0.40 | 0 | 0.0% | 0.0% | — | **8** | 80.0% | **100.0%** | 0.89 |
| *KIR2DS4* | 66 | 56 | 93.3% | 84.8% | 0.89 | 53 | 96.4% | 80.3% | 0.88 | 61 | 95.3% | 92.4% | 0.94 | 56 | 93.3% | 84.8% | 0.89 | 10 | 31.2% | 15.2% | 0.20 | **65** | **97.0%** | **98.5%** | **0.98** |
| *KIR2DS5* | 14 | 9 | 45.0% | 64.3% | 0.53 | 12 | 80.0% | 85.7% | 0.83 | **13** | **100.0%** | **92.9%** | **0.96** | 4 | 66.7% | 28.6% | 0.40 | 8 | 11.8% | 57.1% | 0.20 | 12 | **100.0%** | 85.7% | 0.92 |
| *KIR3DL1* | 66 | 54 | 81.8% | 81.8% | 0.82 | **62** | 93.9% | **93.9%** | **0.94** | **62** | 93.9% | **93.9%** | **0.94** | 54 | **98.2%** | 81.8% | 0.89 | 15 | 21.4% | 22.7% | 0.22 | 61 | 92.4% | 92.4% | 0.92 |
| *KIR3DL2* | 78 | 7 | **100.0%** | 9.0% | 0.16 | 59 | 75.6% | 75.6% | 0.76 | 69 | 93.2% | 88.5% | 0.91 | 71 | 98.6% | 91.0% | **0.95** | 20 | 38.5% | 25.6% | 0.31 | **72** | 90.0% | **92.3%** | 0.91 |
| *KIR3DL3* | 80 | 57 | 86.4% | 71.2% | 0.78 | 42 | 55.3% | 52.5% | 0.54 | 54 | 69.2% | 67.5% | 0.68 | **70** | **89.7%** | **87.5%** | **0.89** | 36 | 46.2% | 45.0% | 0.46 | 64 | 80.0% | 80.0% | 0.80 |
| *KIR3DP1* | 78 | 35 | **97.2%** | 44.9% | 0.61 | 44 | 68.8% | 56.4% | 0.62 | 61 | 87.1% | 78.2% | 0.82 | 63 | 80.8% | 80.8% | 0.81 | 44 | 56.4% | 56.4% | 0.56 | **64** | 88.9% | **82.1%** | **0.85** |
| *KIR3DS1* | 12 | 11 | 16.7% | 91.7% | 0.28 | 10 | 76.9% | 83.3% | 0.80 | 11 | 91.7% | 91.7% | 0.92 | 11 | 78.6% | 91.7% | 0.85 | **12** | 15.0% | **100.0%** | 0.26 | **12** | **100.0%** | **100.0%** | **1.00** |
| All | 749 | 464 | 69.9% | 61.9% | 0.66 | 539 | 74.4% | 72.0% | 0.73 | 647 | 90.6% | 86.4% | 0.88 | 580 | 86.6% | 77.4% | 0.82 | 343 | 31.1% | 45.8% | 0.37 | **682** | **91.4%** | **91.1%** | **0.91** |
